# Brain network dynamics during multi-task demands predict children academic achievement

**DOI:** 10.1101/2025.03.28.645841

**Authors:** Junjie Cui, Rui Chen, Yuyao Zhao, Rui Ding, Sha Tao, Hui Zhao, Shaozheng Qin

## Abstract

Dynamic reconfiguration of neural network and flexible information integration across multiple tasks has been considered critical to characterize individual difference in complex cognition and general intelligence. A promising and underexplored question is how these neurocognitive processes related to children’s academic achievements, a hallmark of high-order cognitive abilities that integrate attention, memory and problem-solving. By using of a multitasking paradigm which bridging outside- in and inside-out approaches, we investigated the dynamic neural mechanisms underlying two core domains of academic performance: math and reading. We first apply partial least squares regression (PLSR) to examine static neural patterns and find that the first latent component—reflecting a generalized brain functional system—predicts math achievement but not reading. The multiple-demand system and the somato-cognitive action network (SCAN) are consistently engaged across diverse task demands. Furthermore, we use a Hidden Markov Model (HMM) to examine dynamic features of brain activity and identify distinct integrated and segregated brain states. Notably, the segregated state—characterized by heightened cortical network segregation—is associated with better math performance. Information-theoretic analyses further reveal that greater complexity in the temporal sequence of the segregated brain functional networks, along with stronger cerebrocerebellar functional coupling, correlates with higher math achievement. By means of multitasks design, these findings suggest that flexible engagement of specialized brain network and automatic information processing is crucial for math learning in children.

## Introduction

With the recent sprout of multitasking paradigms, the task demands emerged from interaction between an individual’s goal, stimuli input and behavioral response is believed to drive neural activity and cognitive flexibility. Therefore, the dynamic and flexible neural circles have been revealed across a variety of tasks was considered to play a crucial role in fundamental mental processes such as integrated intelligence ((Barbey, 2018; Ito et al., 2022; Nau et al., 2024; Shine et al., 2019)). Some computational neuroscientists have shown that the presence of core system in flexible neural networks can enhance performance across tasks—these hubs serve as integrating distributed information (Ito et al., 2022). These components roughly correspond to the action mode network (AMN) identified in neuroimaging analyses. Similarly, the frontoparietal network (FPN) has been shown to coordinate brain-wide activity through flexible reorganization of functional connectivity during multiple tasks (Cole et al., 2013). At the neural level, the multiple-demand (MD) system represents a key component characterized by intensive internal connectivity and consistent activation across diverse tasks. It is believed to facilitate complex problem-solving by integrating subgoals into overarching goals (Duncan, 2013; Duncan et al., 2020). Given its broad task involvement, understanding how these basic neural components support intelligent behavior may reveal core signatures of specific brain function. Interestingly, despite differences in methodological approaches, converging evidence points toward a shared set of specialized, generalized neural mechanisms. The multitasking paradigm thus offers a unique opportunity to assess individual variability in these components from multiple dimensions.

Academic achievement is a multifaceted construct reflecting a range of abilities throughout the entire learning and performance process (Ashkenazi et al., 2013; Floyd et al., 2003; Geary, 2004). It is also a strong predictor of future career success and socioeconomic outcomes (Rivera-Batiz, 1992). Among academic domains, mathematics and reading serve as foundational pillars, shaping many other areas of cognitive and academic development (Ashkenazi et al., 2013; Houdé et al., 2010; Kersey et al., 2019). Prior studies have largely examined how specific mental processes relate to particular academic domains, often overlooking the composite nature of academic performance. Because academic achievement involves multiple cognitive components, capturing its neural basis through single-task paradigms is inherently limited. A more generalized, integrative perspective is thus necessary to better understand the underlying neural architecture. In this context, the inside-out approach provides a framework for moving beyond conceptual constraints and identifying more specialized neural mechanisms (Nau et al., 2024).

Thanks to advances in network neuroscience and neural dynamics, it has become increasingly clear that the human brain meets the demands of an ever-changing world through the flexible and dynamic interaction of different functional networks (Shine, Bissett, et al., 2016; Shine et al., 2019; Shine & Poldrack, 2018). Shine et al. (2019) found that the brain operates within low-dimensional manifolds that are consistent across various tasks, highlighting that the dynamic properties of generalized neural components can account for individual differences in fluid intelligence. It is obvious that dynamic signatures of specific mental processes capture individual differences in some characteristics. Hidden Markov Modeling (HMM) offers a powerful framework for investigating these processes, as it segments observed data into discrete functional states that reoccur over time while simultaneously considering the activity of multiple brain regions. This method has been used to explore the underlying dynamic functional mechanisms and their relationships with behavioral performance (Cai, 2024; Cai et al., 2021; Taghia et al., 2018; Vidaurre et al., 2017).

To verify our hypothesis that generalized neural components can capture a comprehensive profile of neural components underpinning specific academic achievements and to explore the influence of its dynamic signatures on individual differences, we separately applied the same analysis pipeline to two domains, math and reading. At first, we computed the static brain connectivity matrix for each task and applied Partial Least Squares Regression (PLSR) to identify robust and reliable associations between multi-task brain connectivity and academic achievements, which was assessed through a composite test covering a variety of subject-specific contents. After that, we employed a Hidden Markov Model (HMM) to investigate the dynamic reconfiguration of latent neural components and identified six brain states representing distinct cognitive processing modes. In the end, we examined their contribution to academic achievement and further applied graph theory and sample entropy to characterize the spatiotemporal features of these brain states and interpret their associations with performance.

## Results

### The multiple-demand system and somato-cognitive action network associated with math achievement in primary education

Our participants performed multiple tasks in 3T functional magnetic resonance imaging (fMRI) to assess their distinctive basic abilities, involving cognitive, motivational and affective factors (Fig. 1A). We also collected their math and reading achievement test scores out of scanner. Preprocessed BOLD time series were extracted using 268 brain parcels, including the cortex, subcortex, and cerebellum (Fig.1B). Brain connectivity was calculated by Pearson correlation and then transformed into Fisher Z-score. In the end, we obtained four tasks’ brain connectivity matrices, reshaped the upper triangle of each matrix into one row and concatenated them together.

**Figure 1.**
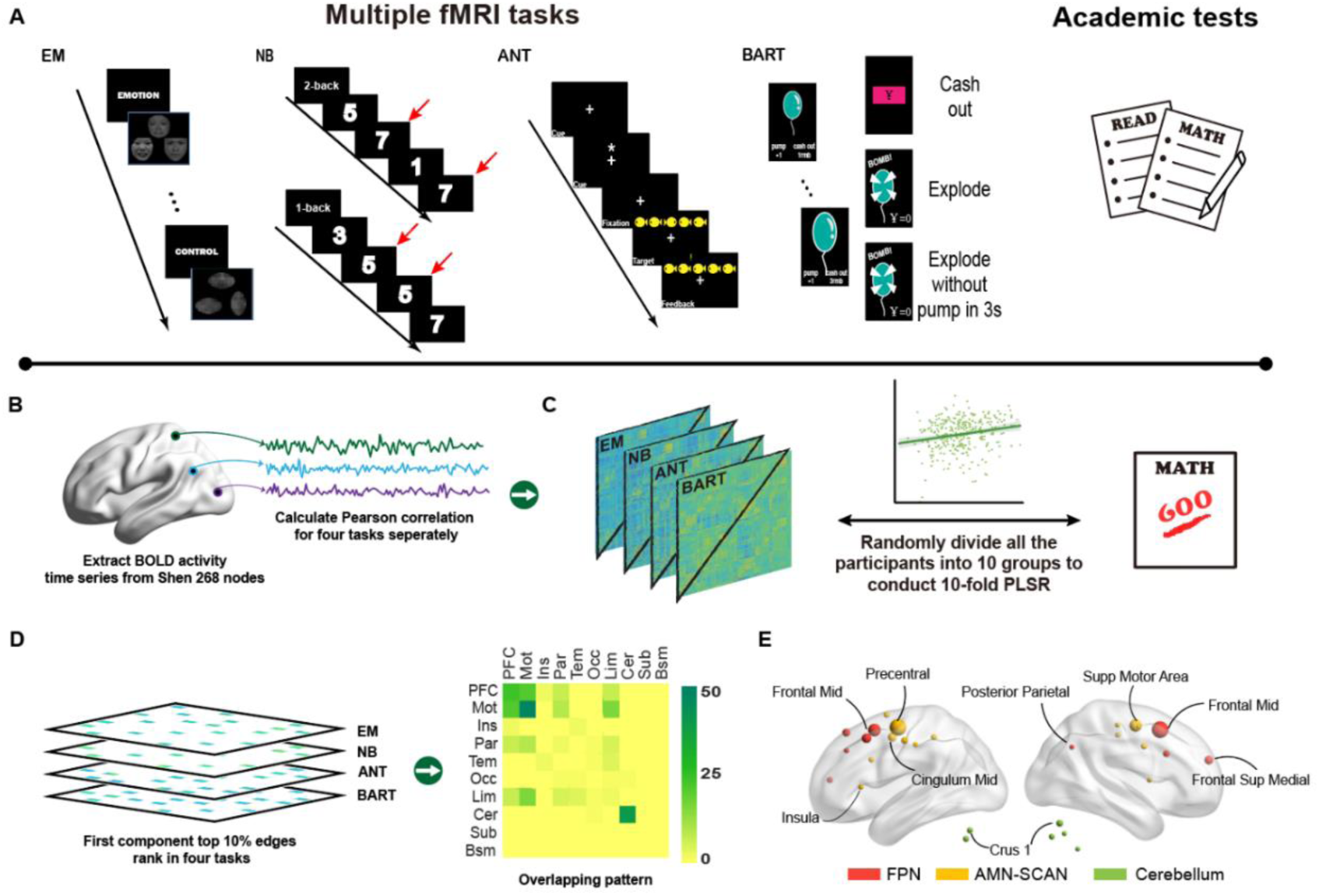
Task designs, static analysis pipeline and partial results. A. Task designs for cognitive, affective and motivational tasks in fMRI, emotion matching task, n-back task, attention network test (children-friendly version) and balloon analog risk task. Out of scanner, children completed mathematic and reading academic achievement tests. B. The average BOLD time series was separately extracted using Shen 268 atlas from tasks. C. Each of brain connectivity matrix was calculated by Pearson correlation and transformed into Z-score. We reshaped the upper triangle of brain connectivity matrix into one row and concatenated them together. 10-fold PLSR was used for identifying the relationship between four tasks and academic performance. D. The process of obtaining the generalized neural components associating with math performance across four tasks. Top 10% was applied as the threshold to select the most stable brain connectivity. E. Illustration of brain regions before the first turning point involved in the first generalized neural component. Size implies the sums of brain connectivity loadings from one brain region, indicating the importance of regions in communication among these regions. Different colors correspond to the following analysis results of Louvain Community Detection, roughly corresponding the canonical brain networks, FPN, AMN-SCAN and cerebellum.

At first, we planned to validate our hypothesis about the association between the specific brain functional system and math cognition. To derive robust results, we applied 100 times 10-fold Partial Least Squares Regression (PLSR) to concatenated brain connectivity and math achievement score, using the average Pearson correlation between predicted math score and actual score as a measure of the relationship between four tasks’ brain activity and math academic performance. Age and gender were regressed from both brain connectivity and math academic performance to avoid the relationship derived by those confounding factors. We adjusted the number of latent components from 1 to 20, with the results of the first component accounting for the largest proportion about their relationship (the latent component = 1, the averaged r = 0.224, permutation p-value = 0.015) (Fig.1C and Fig.S1). For the first component including the rich information about math-relevant neural components across four tasks, we only analyzed the first component in subsequent analyses.

To identify the stable neural pattern associated with math cognition across multiple tasks in all participants, we applied PLSR to all participants in 2000 bootstrap tests with one latent component and obtained the top 10% brain connectivity ranking in four fMRI tasks with the threshold of the bootstrapped loading p-value (-log(P) > 7.358). After restoring the brain connectivity to its original configuration, we obtained four distinct neural patterns of each task, with the n-back task dominating the majority of the selected connectivity (61%) and ranking highest among multiple tasks, suggesting the first component of neural substrates might associate with cognitive control (Fig.S2). Consistent with our hypotheses, we found that there was an overlapping neural pattern across four tasks, comprising 135 functional connectivity and involving 85 brain regions. (Fig.1D, Table.S2). We found that the large proportion of brain connectivity was right located in the MD system and the recently discovered SCAN, encompassing brain regions in the lateral prefrontal cortex, motor cortex, parietal cortex, limbic system, and cerebellum (Duncan, 2010, 2013).

For further exploring the hierarchical signature of the distributed neural pattern, all edge loadings were summed by node, and then we obtained a list of 88 brain regions’ ranking scores. We found that the top 30 brain regions before the first turning point roughly overlapped with the MD system and the SCAN (Fig.1E, Fig.S3, Table.S2), suggesting that the specific component with intense inner-connectivity across multiple tasks might serve as a core system for integrating information from distributed brain regions, consistent with previous electrophysiological evidence for binding of cognitive operating to their contents (Duncan, 2013; Duncan et al., 2020).

### Math-relevant brain states share cerebral functional modules but differ in cerebrocerebellar functional coupling

With the belief that the brain operates in a highly dynamic way, we were interested in the dynamic signature of the math-specific neural component operating processes and its contribution to math cognition, further disclosing the characteristics of math cognition in primary education. Previous study demonstrated a low-dimensional manifold existed in whole-brain dynamics across multiple tasks, with specific components playing a general role, separately associated with the integrated and segregated states (Shine et al., 2019). Thus, we hypothesize that the specific neural components operate similarly across four tasks, with certain dynamic signatures potentially offering insights into math cognition. To test our hypothesis, we extracted the BOLD time series of 30 brain regions with the maximum sum of loadings identified above, concatenated them together in four tasks, and utilized Hidden Markov Model (HMM) to study its dynamic operating processes (Fig. 2A, 2B). Due to consideration of both the operating process generality across all participants and the nuance of dynamic signature, we used 6 states to fit the long concatenated BOLD time series. The outputs of HMM encompass the activation and connectivity pattern of each state at group level, the probability distribution of each state in every time point, Viterbi path (the most probable decoded state sequence), and transition probability matrix (Fig.2C, 2D, Fig.S4).

**Figure 2.**
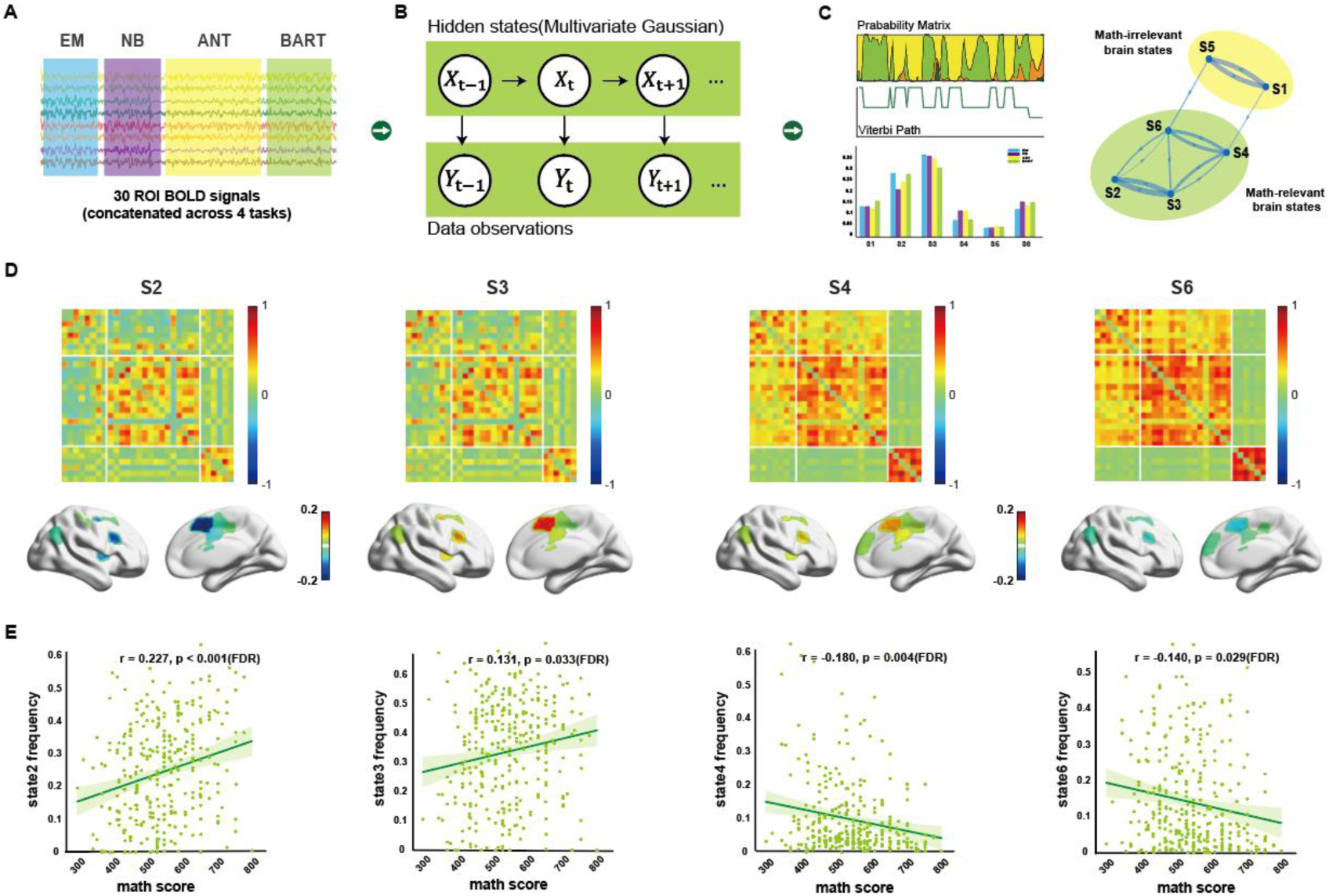
Dynamic analysis pipeline and the output of HMM. A. The input of HMM. Time series of selected brain regions from four fMRI tasks were concatenated together across all the participants. B. HMM modelling processes. We use multivariate Gaussian distribution to fit each brain state. C. The output of HMM. The distribution probability of each brain state in every time point (Left upper). Viterbi path, the most possible brain state sequence decoded for every participant (Left middle). The frequency of each brain state among different tasks (Left bottom). Transition probability in group level (tp>0.1) (Right). D. Connectivity pattern and activation pattern of four task-dependent states at the group level. E. The Pearson correlation between math academic performance and brain states’ frequency in four tasks. All the correlations were FDR-corrected among six states.

We observed a positive correlation between the frequency of both s2 and s3 and math achievement (Fig. 2E. s2: r = 0.227, p-value < 0.001; s3: r = 0.131, p-value = 0.033, FDR-corrected), with s4 and s6 showing a negative correlation (Fig. 2E. s4: r = -0180, p-value = 0.004; s6: r = 0.140, p-value = 0.029, FDR-corrected). We applied the Louvain Community Detection algorithm to the transition probability matrix to explore their temporal hierarchical signature and found State 2 (s2) and State 3 (s3), State 4 (s4) and State 6 (s6), State 1 (s1) and State 5 (s5) can be grouped into three distinct meta-states, that the brain tends to cycle within and are commonly associated with different cognitive processes (Shine, Koyejo, et al., 2016; Vidaurre et al., 2017). The correlation results can be further explained by their different spatial organization patterns. To further understand their distinct function in operating processes, we applied the Louvain Community Detection algorithm to states’ functional connectivity pattern at the group level. Consistent with previous studies, those regions can be grouped into three modules, roughly corresponding to FPN, AMN-SCAN, and cerebellar regions (Crittenden et al., 2016; Duncan, 2013)(Table.S5). We observed that math-relevant brain states share same cerebral functional modules, while different meta-states exhibit distinct patterns of cerebro-cerebellar functional coupling. In states s2 and s3, non-motor regions of the cerebellum—specifically Crus I, Crus II, and lobule IX—show stronger link with the frontoparietal network (FPN).

Compared with four math-relevant states, another two states show a minimal difference in cerebral organization. The left insula and right supplementary motor area (SMA), supposed involved in AMN-SCAN, are assigned to the FPN. In previous research, the insula was regarded as a potential candidate hub for network switching signaling (Cai et al., 2021; Molnar-Szakacs & Uddin, 2022), and that can be a reasonable explanation why two brain states are not relevant to math cognition, for they are not engaged in working processes but might function at the indispensable intermediate stages and play a crucial role in switching among different organizations.

### The functionally specialized segregated brain states contribute to math achievement

To describe the spatiotemporal characteristics of the specific neural components in a more nuanced way, we adopted concepts from information theory and graph theory to explore the temporal and spatial signature of each brain state. Sample entropy is a measure used to quantify the complexity of biological signal, where a higher value indicates greater unpredictability and a higher capacity for information processing. So here we want to explore whether the neural entropy of different brain states is related to math cognition, identifying some brain states capacity of information processing essential to math performance. After controlling age and gender, neural entropy of both s2 and s3 was still positively correlated with math academic performance (s2: r = 0.214, p < 0.001; s3: r = 0.166, p = 0.01, FDR-corrected. Fig. 3A), while other two task-dependent states negatively related (s4: r = -0.160, p = 0.010; s6: r = -0.122, p = 0.048, FDR-corrected. Fig. 3A). Again, we didn’t find any relationship between task-independent states and math cognition.

**Figure 3.**
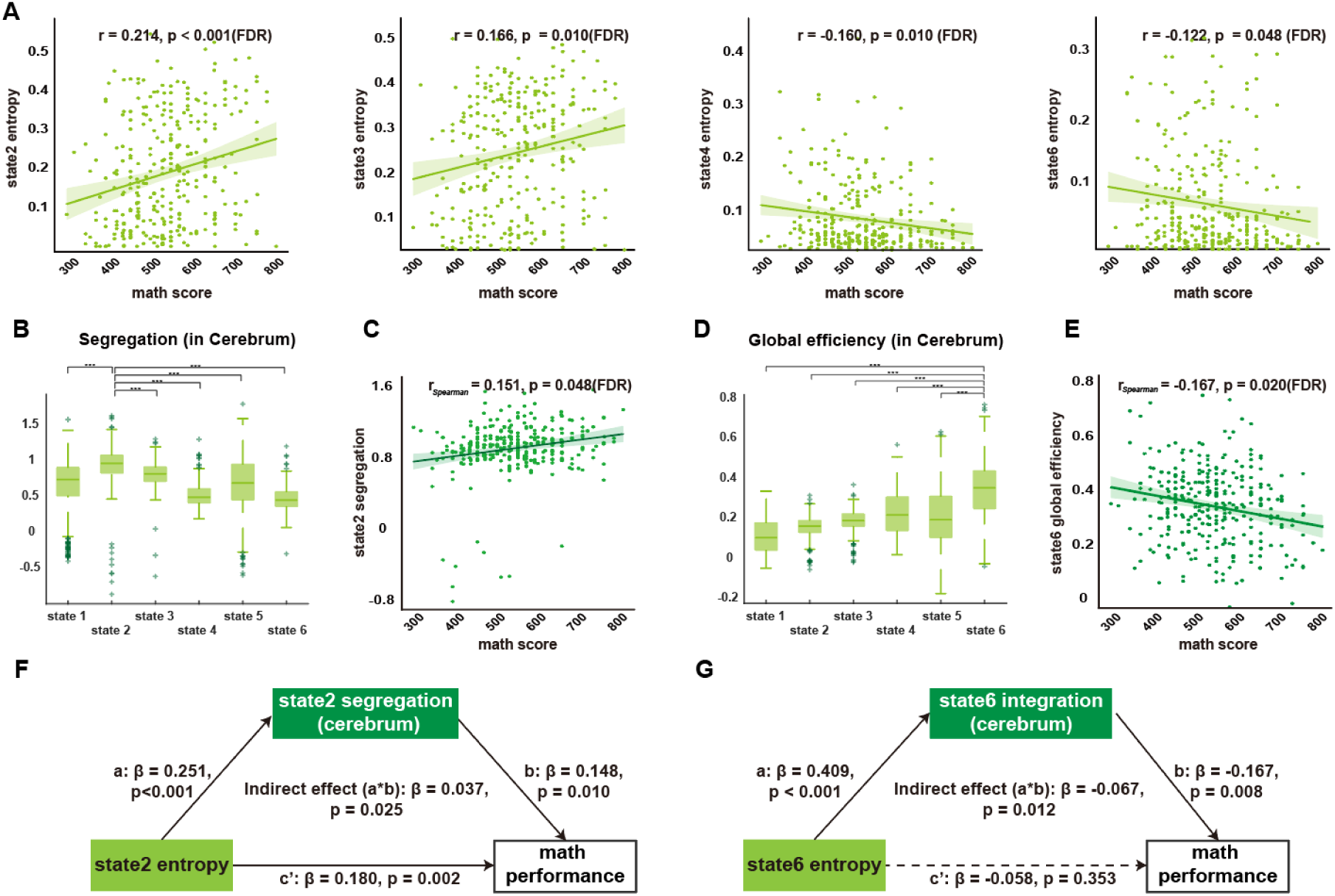
The spatiotemporal signature of each brain state. A. The Pearson correlation between math academic performance and neural entropy of probability sequence of each brain state (FDR-corrected). B. The segregation of cerebral cortex parts in each brain state, with the results showing that s2 is highest among six states. C. Spearman correlation between math academic performance and s2 segregation (FDR-corrected). D. The global efficiency of cerebral cortex parts in each brain state. s6 is highest among six states. E. Spearman correlation between math academic performance and s6 global efficiency (FDR-corrected). F.G. The mediation analysis of state entropy, state spatial features, and math academic performance.

To further investigate what leads to the distinct relationships between the temporal features of each brain state and math academic achievement, we utilized function in HMM-MAR toolbox to re-infer brain states at the individual level and applied graph theory to explore their topological signature, aiming to uncover the underlying causes. For the integration and segregation manifolds in human brain dynamics have been demonstrated for decades of years, both are associated with distinct cognitive processes and contribute to behavior performance (Capouskova et al., 2023; Cohen & D’Esposito, 2016; Shine, Bissett, et al., 2016; Shine & Poldrack, 2018). For the hierarchical nature of human brain, here we utilized network segregation and global efficiency to quantify segregation and integration of cerebral parts in each brain state. Segregation quantifies the difference between the average within-network connectivity and the average between-network connectivity (Chan et al., 2014). Owing to diverse individual differences, we used the Louvain Community Detection algorithm to identify the network community for each participant and applied network segregation to their individual community (Gratton et al., 2018). The global efficiency is the average inverse shortest path length in the network, with a higher measure indicating an intensive integration. We found that s2 is the most segregated state, with a positive correlation with math academic performance uniquely (*r*_*spearman*_ = 0.151, p = 0.048, FDR-corrected. Fig. 3B, 3C); whilst the s6, with the highest global efficiency, showed the only negative correlation with math academic performance (*r*_*spearman*_ = 0.167, p = 0.020, FDR-corrected. Fig. 3D, 3E). It seems that the segregated cerebral neural components contributes to math academic performance, which suggested that s2 may serve as an expertness state and render brain networks functionally specialized (Sporns & Betzel, 2016; Wig, 2017).

To the best of our knowledge, the emergence of diverse brain functions results from the spatiotemporal interactions within distinct brain organizations. To test our hypothesis that the varying associations between the temporal features of each state and math cognition are attributable to their unique spatial organization, we performed several mediation analyses using structural equation modeling (SEM). Our results revealed that the segregation of s2 mediated the relationship between the neural entropy of s2 and math academic performance (indirect effect (a*b) = 0.037, p = 0.025; Fig. 3F), whilst the global efficiency of s6 mediated the relationship between the neural entropy of s6 and math academic performance (indirect effect (a*b) = -0.067, p = 0.012; Fig. 3G). The results suggest that it is exactly the distinct spatial organization of each brain state drive the differential relationships between their temporal features and math academic performance.

### Cerebrocerebellar Coupling Plays a Crucial Role in Math Cognition

Having validated that s2 with the most segregated cerebral cortex promotes math cognition, we proposed that s2 represents an efficient expert state characterized by functionally specialized brain community. The coordinated organization not only embodied in the organized cerebral cortex but also in the connections between the reciprocal regions in the cerebrum and cerebellum. Recently, the cerebellum was considered to facilitate human cognition such as cognitive control but not limited to motor coordination. Many studies indicated that there are some counterpart organizations in the cerebellum corresponding to the cerebral cortex and demonstrate cerebrocerebellar functional coupling benefits learning and task performance (Balsters & Ramnani, 2011; Boven et al., 2023; Clark et al., 2022). As a result, here we hypothesized that the reciprocal cerebrocerebellar functional coupling might be essential to math cognition.

To test our hypothesis, we defined an index measuring cerebrocerebellar functional coupling in the math-specific neural components and applied one-way ANOVA analysis to six states, finding that s2 exhibits the top coupling among six states, followed by s3 (Fig. 4A). Both brain states contribute to math achievement performance. To study the distinctive functional coupling organizations in s2 more detailed, we utilized paired t-test to compare the brain connectivity of s2 with its counterpart (s6) in another working meta-states, identifying the stronger brain connections. All contrasts were FDR-corrected. As illustrated in Fig. 4B, we observed a distinct coupling pattern between motor regions and non-motor regions in the cerebellum. Compared to s6, motor regions in the cerebellum have stronger connections with regions in CON and sensorimotor cortex in s2, whereas non-motor regions are strongly linked to higher-order brain regions in FPN. Considering the series of positive correlations between s2 and math cognition, we believe that cerebrocerebellar coupling contributes to math cognition as well.

**Figure 4.**
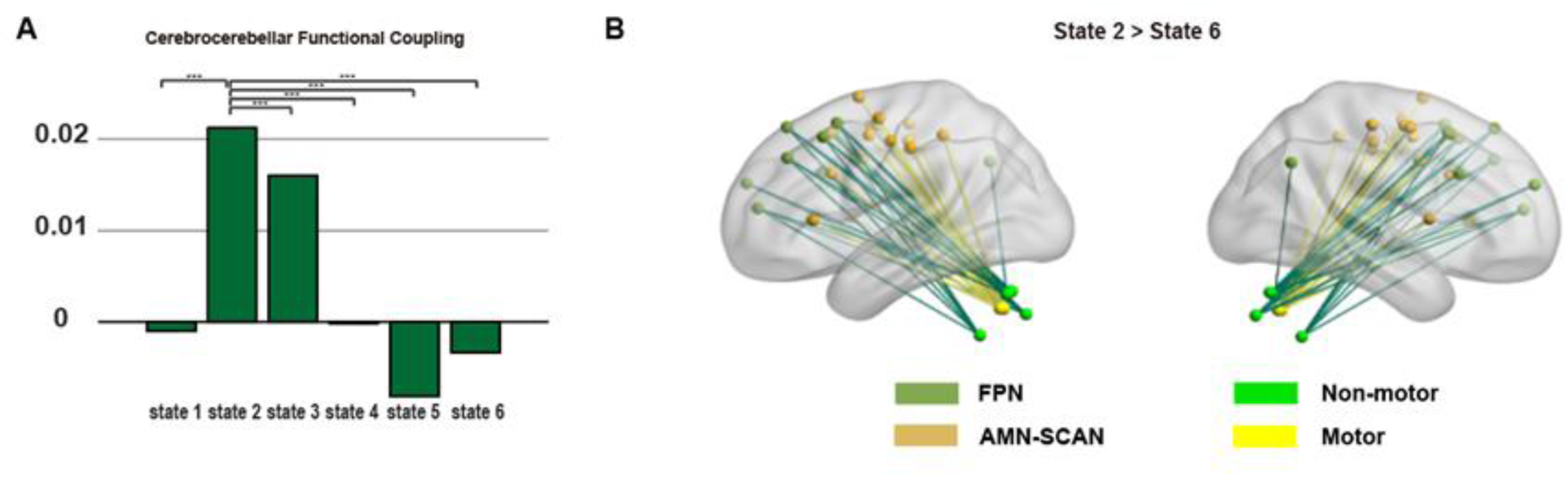
Cerebrocerebellar functional coupling measure of each brain state and visualization. A. Cerebrocerebellar coupling index. The higher index indicates a stronger coupling between the reciprocal parts in the cerebral cortex and cerebellum. B. The visualization of stronger brain connectivity between cerebrum and cerebellum in state2 than state6.

### Reading analysis

We have applied the same analysis pipeline to reading performance and found that the predictive pattern is different from math, that no one generalized neural pattern can predict reading performance. After controlling age and gender, the first component is not enough to predict reading performance (Pearson r = 0.106, permutated p-value = 0.079, details in Fig. S5). We also associated the frequency of brain states of math-relevant brain states with reading performance, there is no significant correlation. So we proposed that MD system and SCAN might play a specific role in math cognition, but not general neural substrates in academic performance, which further disclose the particular characteristics of math performance.

## Discussion

Our results revealed the association between the MD system, SCAN and math cognition from the dynamic perspective, offering insights into the neural mechanism of math cognition during primary school years. Consistent with our hypotheses, we identified one generalized neural component relating to math cognition across cognitive, affective and motivational tasks, encompassing the MD system characterized by intense inner-connectivity. We further investigated the dynamic operating processes of math-related generalized neural component using HMM, which highlighted the interplay between integrated and segregated states. We also found the segregated state exhibiting heightened cortical network segregation—is associated with better math performance.

Information-theoretic analyses further reveal that greater temporal complexity of the segregated brain functional networks, along with stronger cerebrocerebellar functional coupling, correlates with higher math achievement. These findings suggest that flexible engagement of specialized and automatic information processing is crucial for early math learning. Contrast to math performance, we did not observe a similar neural pattern in reading performance, further validating the dissociable neural components underlying math and reading performance.

It was the first study that identified the association between the MD system and SCAN in non-math tasks and math cognition, demonstrating that such neural components supporting multiple task demands are especially required during math problem solving. Previous research has identified a ‘math-responsive brain network’, which is activated when participants solve various types of simple math problems, and proposed that it is a math-specific brain network, distinct from the language-related brain network (Amalric & Dehaene, 2016, 2019). Our results provided complementary insights by highlighting its association in non-math tasks, which suggest a unique demand for cognitive control and information integration in primary math cognition. It has been observed that compared with adult participants, child participants show a higher activation in the frontal cortex when solving arithmetic problems (Zelazo & Carlson, 2020). The term ‘multiple-demand’ system is attributed to its ubiquitous activation among multiple tasks, thought to play a crucial role in general cognitive control during goal-directed tasks (Duncan, 2013; Duncan et al., 2020). Aligning with our results, working memory task accounted for the majority (52%) of the selected brain connectivity among the four tasks. To the best of our knowledge, working memory exhibits the highest demand for cognitive control among those tasks. In one recent research, researcher explored the controllability of brain structural network from a perspective of control theory. They validated that the modal controllability of MD system is ranked high, which means those brain regions can push the brain into difficult-to-reach states requiring substantial input energy (Gu et al., 2015). Beyond cognitive control theory, the MD system has been considered a potential neural substrate for integrated intelligence, capable of integrating information from distributed brain regions, due to its strong inner connectivity in contrast to outer brain networks (Assem et al., 2020; Duncan et al., 2020). In primary math education, procedural skills are essential for solving arithmetic problems, requiring integrated intelligence. Previous research has shown that fluid intelligence is an effective predictor of math achievements, which may help explain the relationship between two (Assem et al., 2020; Green et al., 2017).

Beyond locating the related neural substrates, we demonstrated the underlying operating processes of the MD system in terms of integrated and segregated manifolds for the first time. Previous research has demonstrated that the whole brain flexibly switches between segregated and integrated states to meet task demands in a time-sensitive manner, beneficial for efficient information processing and cognitive flexibility (Capouskova et al., 2023; Shine, Bissett, et al., 2016; Shine et al., 2019). Actually, MD system corresponds to large-scale resting-state brain networks in anatomy, FPN and AMN. While MD system was often conceptualized as operating as a unified system from a relatively static perspective, its dynamic aspects have been largely overlooked (Crittenden et al., 2016). To further characterize the role of the MD system in math cognition, we investigated its dynamic functioning across four tasks, aiming to identify generalizable neural mechanisms and promote their application in more naturalistic contexts (Nau et al., 2024). Consistent with previous results, MD system exhibited both integrated and segregated states, which likely play distinct roles in task-specific demands and contribute to math performance. Additionally, we observed evidence of intermediate brain states that may serve as transitional phases. This was supported by Louvain community detection results, which revealed that the insula was reassigned to a different community, specifically the FPN. Anterior insula was considered as a gatekeeper of executive control, which plays a role in orchestrating and driving activity of other major functional brain networks, such as FPN (Molnar-Szakacs & Uddin, 2022).

The segregated MD system contributes to math performance during primary education, with stronger connectivity between reciprocal brain regions in the cerebral cortex and cerebellum. Segregated brain systems are commonly associated with specialized and expert information processing, which facilitates task demands (Baum et al., 2017; Capouskova et al., 2023; Wig, 2017). Previous research has shown that, with practice, brain states exhibit greater segregation, reflecting the development of automatic behaviors (Bassett et al., 2011, 2015). In the context of math cognition, children with greater procedural skill capacity require less executive function, relying on more automatic processes (Cragg et al., 2017). In this brain state, stronger reciprocal connectivity between the cortex and cerebellum supports more efficient information processing. Prior studies have demonstrated that cerebellum-cingulo-opercular network connectivity strengthens during childhood, enhancing attention efficiency (Clark et al., 2022). These findings collectively highlight the role of practice in primary school math education. These results align with the neural efficiency theory, which suggests that individuals exhibit more efficient brain activity during cognitive tasks, offering additional insights into the function of multiple-demand systems from the perspective of neural dynamics (Neubauer & Fink, 2009). Consistent with previous research, segregated brain states with higher entropy have been shown to play a crucial in cognitive flexibility, enabling rapid adaptation to task demands.

MD system and SCAN play a specific role in math cognition, but not a domain-general role in academic performance. To investigate this, the same analysis pipeline was applied to reading performance. Surprisingly, we didn ‘ t observe similar results, consistent with substantial evidence indicating distinct neurocognitive mechanism for reading and math cognition. Prior studies have demonstrated that fluid intelligence is supported by the MD system rather than the language system (Blanke et al., 2014; Diachek et al., 2020; Fedorenko et al., 2024). Furthermore, research enhancing the difficulty of sentence comprehension has shown that language and math rely on separate neurocognitive systems. These findings support our hypothesis that MD systems serve as a specialized neural substrate underpinning math cognition, highlighting the cognitive control demands associated with primary school math education.

Our research has some limitations. First, we didn’t consider the interaction between the generalized neural component with other brain networks, such as default mode network, for the brain often functions as an integrated whole to adaptively respond to real-world challenges. Future research could investigate its operating processes from a whole-brain perspective to uncover additional mechanisms underlying it. Second, while we employed multiple laboratory tasks to explore the dynamic operating processes of the generalized neural component, hoping to uncover its mechanism from a more generalized perspective, the results still lack broader generalizability. It would be valuable to examine its dynamic nature in more naturalistic contexts to enhance ecological validity. Third, we may add more academic fMRI tasks to extend its generality and increase ecological validity. In conclusion, our findings demonstrate that the MD system and SCAN play a specific role in primary math cognition, with its operating processes adhering to the principles of integrated and segregated manifolds. Furthermore, in primary school, higher math achievement is associated with more flexibly specialized information processing.

## Materials and methods

### Participants

In math cognition analysis, 310 participants (ranging from 6 to 12 years old, 52.6% female) were included in analysis with completely cognitive and affective tasks imaging data and math academic test, which were derived from the Children school functions and Brain Development Project (CBD, Beijing Cohort). All participants reported no history of vision problems, no history of neurological or psychiatric disorders, and no current use of any medication or recreational drugs. Written informed consent was given by each child and one of their parents or legal guardians. The procedures of consent and experiment were approved by the local Ethics Committee and were in accordance with the standards of the Declaration of Helsinki. In the control analysis, reading performance was matched for each participant included in the math cognition analysis. After matching, 305 students with available reading performance were included. The demographics of the dataset for two analyses are listed in Table S1.

### Image data acquisition

Brain imaging data were obtained by the same type of Siemens 3.0T scanner (Magnetom Prisma syngo MR D13D, Erlangen, Germany) with a 64-channel head coil and T2*-sensitive echo-planar imaging (EPI) sequence based on blood oxygenation level-dependent contrast. Thirty-three axial slices (3.5 mm thick, 0.7 mm skip) parallel to the anterior and posterior commissural line and covering the whole brain were imaged with the following parameters: repetition time (TR) = 2000 ms, echo time (TE) = 30 ms, flip angle = 90°; voxel size = 3.5 × 3.5 × 3.5 *mm*^3^, and field of view (FOV) = 224 × 224 *mm*^2^ . In addition, high-resolution anatomical images from each participant were acquired by three-dimensional sagittal T1-weighted magnetization-prepared rapid gradient echo (MPRAGE) with a total of 192 slices (TR = 2530 ms, TE = 2.98 ms, flip angle = 7°, inversion time = 1100ms, voxel size = 1.0 × 1.0 × 1.0 *mm*^3^, acquisition matrix = 256 × 224, FOV = 256 × 224 *mm*^2^, brand width = 240 Hz/Px, slice thickness = 1 mm).

### Image data preprocessing

Brain images were preprocessed with the fMRIPrep 1.4.1 (Esteban et al., 2019) pipeline implemented in Nipype 1.2.0 (Gorgolewski et al., 2011). The first 4 volumes of each run for all tasks were discarded for signal equilibrium and the adaptation of participants to fMRI scanning noise. For each participant, the following preprocessing procedures were conducted. First, a reference volume and its skull-stripped version were generated using a custom methodology of fMRIPrep. Co-registration was configured with nine degrees of freedom to account for distortions remaining in the BOLD reference. Head-motion parameters with respect to the BOLD reference (transformation matrices and six corresponding rotation and translation parameters) were estimated before any spatiotemporal filtering using mcflirt (FSL 5.0.9). Functional images of each task were slice-time corrected using 3dTshift from AFNI. The resultant images (including slice-timing correction when applied) were resampled into their original, native space by applying a single composite transformation to correct for head-motion and susceptibility distortions. These resampled images are referred to as preprocessed BOLD functional images in the original space. After that, images were resampled into a standard space, generating a preprocessed BOLD run in the ‘MNI152NLin6Asym’ space. For more robust research results, participants with excessive head motions (more than 1/3 frames with standardized DVARS >1.5 or frame displacement > 0.5) or incomplete scanning were excluded from further analyses. After obtaining the above preprocessed data, we further regressed out 24 head motion parameters, white matter, cerebrospinal fluid (CSF), and global signal as confounding factors. The preprocessed BOLD signals were then detrended to remove low-frequency drifts and linear trends that could arise from scanner noise or physiological artifacts. A bandpass filter was then applied, restricting the frequency range of the data to 0.009-0.015 Hz.

### Brain parcellation and brain connectivity matrices

After preprocessing, we extracted the average BOLD time series from 268 predefined regions of interest. The parcellation was generated using a group-level spectral clustering algorithm to applied to an independent dataset. Based on their location in the brain, the 268 nodes were assigned to ten canonical brain networks: PFC, Mot, Ins, Par, Tem, Occ, Lim, Cer, Sub, Bsm. The size of brain connectivity matrix is 268*268, with each brain connectivity calculated through Pearson correlation and transformed into Fisher z-score in GRETNA toolbox ([NITRC: GRETNA: Tool/Resource Info](https://www.nitrc.org/projects/gretna/)).

### Math and reading academic tests

#### Mathematics achievement test

The mathematics achievement test used in this study was developed by the National Children’s Study of China (NCSC) in line with Chinese National Curriculum System (Dong and Lin, 2011). It is a comprehensive academic achievement test that assesses arithmetic, geometry, and problem-solving skills through questions aligned with grade-level standards. Item response theory (IRT) was used to ensure that scores were comparable across grades. The test was validated on a nationally representative sample of 140,000 children in 600 schools in 100 counties in 31 provinces in mainland China. The Cronbach’s alpha of the test across different grades ranged from 0.83 to 0.89. The test took approximately 45 minutes to complete.

#### Reading achievement test

The reading achievement test, also developed by the NCSC based on the Chinese National Curriculum System (Dong and Lin, 2011), assessed pinyin, character and word recognition, grammar, and comprehension of sentences and passages. The test was tailored to grade-level standards, and IRT was used to ensure cross-grade comparability. The test has been validated on the same nationally representative sample, with Cronbach’s alpha across different grades ranging from 0.81 to 0.89. The test took approximately 45 minutes to complete.

### fMRI cognitive and affective tasks

Before performing tasks in fMRI, every participant was instructed by specialized researcher. In fMRI, all participants completed four tasks in fMRI in the following order, emotion matching task, attention task, working memory task and risk-taking task.

#### Emotion matching task

was a random block experiment design, encompassing emotion matching condition and sensorimotor control condition. During the emotion matching block, participants watched a trio of faces with fear or angry expressions, and selected one of two faces below expressing the same emotion with the target one. While on the sensorimotor control condition, students viewed a trio of geometric shapes and chose one with shape same with the target one. Each block started with a cue for 5s to inform which condition it is now, followed by six trials with 5s each.

#### Attention network test task

was a children-friendly version revised from attention network task. It is a random block design and consists of two factors, the types of cue and flanker. Each trial started from one of the four cue conditions (1) no-cue, (2) double-cue, (3) center-cue, or (4) spatial cue, either above or below the fixation cross at the screen center. The centrally presented target ‘fish’ was flanked by one of the three different types of ‘fish’ stimulus: (1) congruent flanker ‘fish’, (2) incongruent flanker ‘fish’, or (3) only a ‘fish’. **Each trial started with a fixation cross at the central of the screen for 400 ms to 1000 ms. Thereafter, in some of the next trials a warning cue appeared for 150 ms. And a stationary fixation phase of 450 ms was presented after the end of the cue. Thereafter, the target ‘fish’ stimulus with one of two types of flanker (congruent or incongruent) was presented until the participant made a button press or reached the time limit of 1000 ms. The duration of the last fixation was 1000 ms minus the corresponding reaction time. After responding, the participant received visual feedback from the computer. For correct responses, a simple animation sequence showed the target fish blowing bubbles. Incorrect responses had no animation of the fish.**

#### Working memory task

consists of three conditions (0-, 1- and 2-back), with each condition including 4 blocks. One block spanned 27 s, with a random sequence of 15 digits presented to the participant. Each digit was presented for 400 ms, followed by an inter-stimulus interval of 1400 ms. During the 0-back, participants were instructed to identify whether the digit is a “1” or not. During the 2-back, participants were instructed to identify whether the current digit is same with the two positions back one.

#### Risk-taking task

is an event-related design to explore the reward and incentive mechanism. There are three conditions according to the operation of each participant, pump, cash-out, and explode. In each trial, participants can choose to press the ‘pump’ button, or stop by pressing the ‘cash-out’ button within 3000 ms. Otherwise, balloon would explode automatically if there was no response. In each trial, rewards can be accumulated by every pump (1 ¥ per pump) and be added to a total number. However, money would be lost if the balloon exploded within each trial. The number of trials depend on the performance of each participant with a self-paced design.

In sum, EM task spanned 359 s involving 179 volumes; for the WM 462 s and 232 volumes; for the ANT 254 s and 177 volumes; for the BART task 368 s and 184 volumes.

### Partial least squares regression

Before applying PLSR, we reshaped the upper triangle of every task’ connectivity matric into one row and concatenated four tasks’ connectivity together. To avoid the effect of confounding factors, we regressed age and gender from the math academic performance and brain connectivity separately and used their residuals for further analysis.

To validate the solid association of multiple-demand systems and math cognition and avoid overfitting, we adjusted the latent components from 1 to 20 and conducted 10-fold PLSR 100 times under each condition to measure their relationship. As the parameter increases to 20, the average correlation gradually converges to 0.267, permutation p-value < 0.001. However, when the number of latent components equals 1, the average relationship = 0.224, permutation p-value = 0.015. For considering computational efficiency and maximum information content, we only analyzed the first component in the subsequent analysis.

After identifying their relationship, we bootstrapped participants 2000 times and applied PLSR to them to identify the math-related neural pattern across four tasks. We calculated each edge’s z-score of loadings in the first latent component and chose the top 10% edges ranking in four tasks to find the neural pattern underpinning math cognition across cognitive tasks. Through these methods, we can get the most stable and the most math-related neural pattern.

### Hidden Markov Model

To delve into the dynamics of the multiple-demand systems, we applied a Gaussian HMM to the concatenated time series from 30 ROIs which were identified in the static analysis and estimated 6 recurring discrete brain states, implemented by the MATLAB toolbox HMM-MAR (https://github.com/OHBA-analysis/HMM-MAR). Each state was estimated as a multivariate Gaussian normal distribution with first and second order statistics (mean activity and covariance matrix). Actually, we also implemented 4 and 8 states to fit these brain data, but the former cannot seize the nuance of the dynamic processes and the latter included some states specific to part of participants. The output of HMM encompassed both brain activity and connectivity pattern of each state at the group level, the probability time series of each state, Viterbi path, and transition probability matrix. After we obtained all the results at the group level, we used the function in HMM-MAR toolbox to re-infer each state at individual level.

### Sample Entropy

Sample entropy is an index measuring the complexity of physiological signals, and quantifies its degree of unpredictability. After identifying the embedding dimension m, it will count all the number B of similar series clips m and contrast it with the number A of the embedding dimension being m+1. If the time series is very complex, the longer embedding dimension would lead to fewer similar series clips. The larger sample entropy corresponds to the more complex series. Considering our time points is no more than 1000, we obeyed the traditional rules and selected 2 as our embedding dimension with the tolerance being 2*STD.

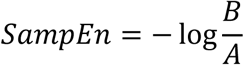

### Graph theory

#### Louvain Community detection

To identify how these states are hierarchically organized in time and community assignments in space, we used Louvain community detection algorithm in Brain Connectivity Toolbox (BCT, https://sites.google.com/site/bctnet/ ) to detect modular partitions. For robust conclusions, every time we applied this algorithm 500 times and used community consensus algorithm to derive the latent structures.

#### Network segregation

Network segregation measures the average brain connectivity difference between inter-networks and inner-networks. For calculating this index, we used the consensus community partition derived in Louvain Community Detection and applied the freely available MATLAB code from https://github.com/mychan24/system_matrix_tools.

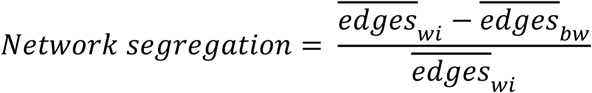

#### Global efficiency

The global efficiency is the average inverse shortest path length in the network. The high global efficiency indicates an intense integration facilitating transfer of information between different nodes.

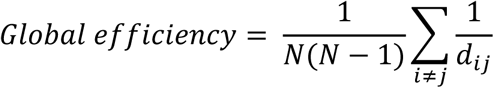

### Mediation Model

We use Mplus Version 8.3 to infer their mediation effect among neural entropy, network segregation, global efficiency and math academic performance. Firstly, we calculated Pearson correlation between the different features of brain states and math academic performance, and all the correlation was FDR-corrected. We found the particularity of s2 and s6 in math cognition. Controlling age and sex as confounding factors, we conducted the mediation analysis among them. The fitness of each model was assessed using a χ2 test, which showed no significance. In addition, the Root Mean Square Error of Approximation (RMSEA) was found to be below 0.08, the Standardized Root Mean Square Residual (SRMR) had an outcome value of less than 0.08, and the Comparative Fit Index (CFI) was above 0.90.

### Cerebrocerebellar Functional Coupling Metric

To quantify the degree of cerebrocerebellar coupling, we defined an index measuring the average brain connectivity difference between Motor areas and non-motor areas in cerebellum to brain network in the cerebral cortex. A higher index (above zero) indicates a more robust cerebrocerebellar coupling for the stronger connectivity exists between the reciprocal networks.

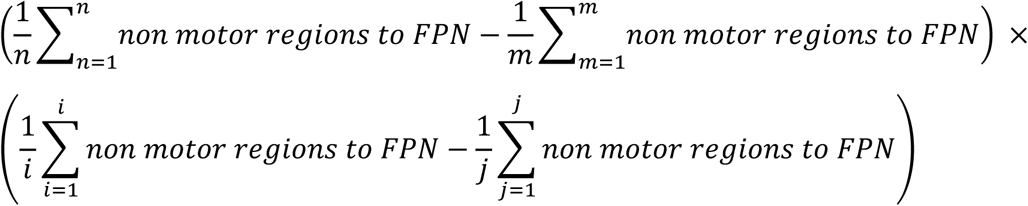

