## Supplemental files for "Brain network dynamics during multi-task demands predict children academic achievement"

**Supplementary figures**

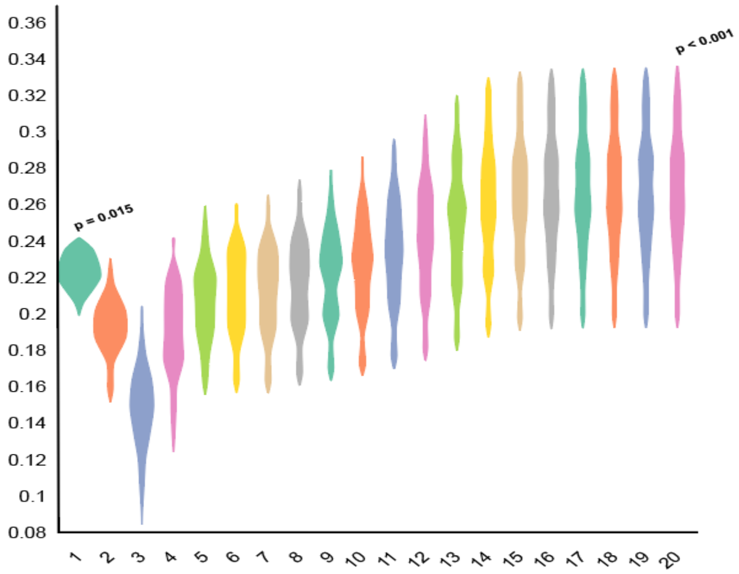


***Figure S1. The adjustments of parameter components in PLSR (Math achievement).*** *To ensure more reliable results, we applied 10-fold PLSR to randomly divided 310 participants 100 times, adjusting the number of latent components from 1 to 20. The average correlation was used to assess the relationship, with permutation testing (2000 iterations) employed to calculate the p-value. Components number = 20, p-value < 0.001.*

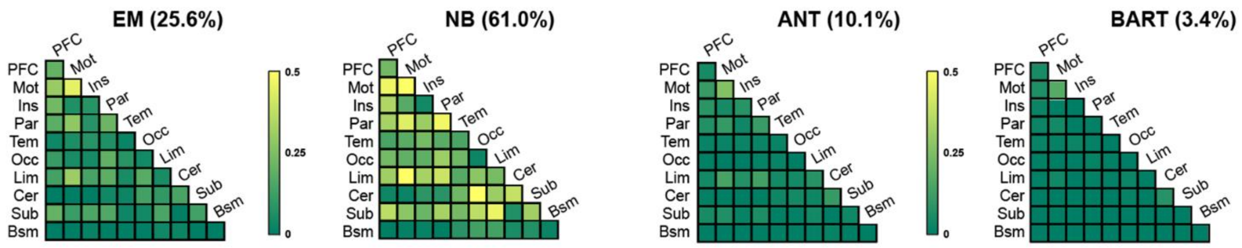


***Figure S2. Different proportions of brain connectivity across four tasks.*** *The N-back task contributes to the majority of the selected connectivity (61%), ranking highest among the four tasks. For illustration, regions were grouped into ten brain modules.*

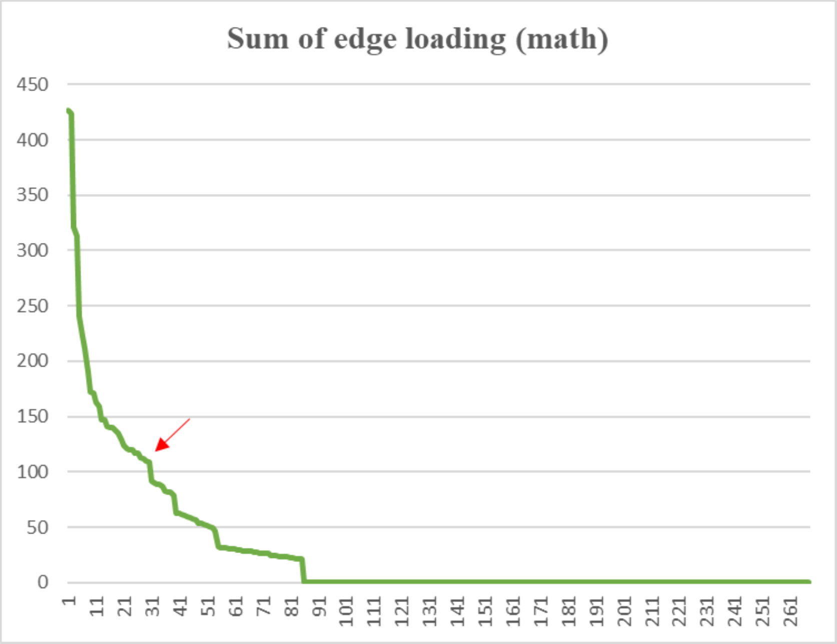


***Figure S3.*** *The ranking of brain regions based on the sum of loadings derived from 2000 bootstraps. The red arrow indicates the first turning point (the 30$^{th}$ brain region).*

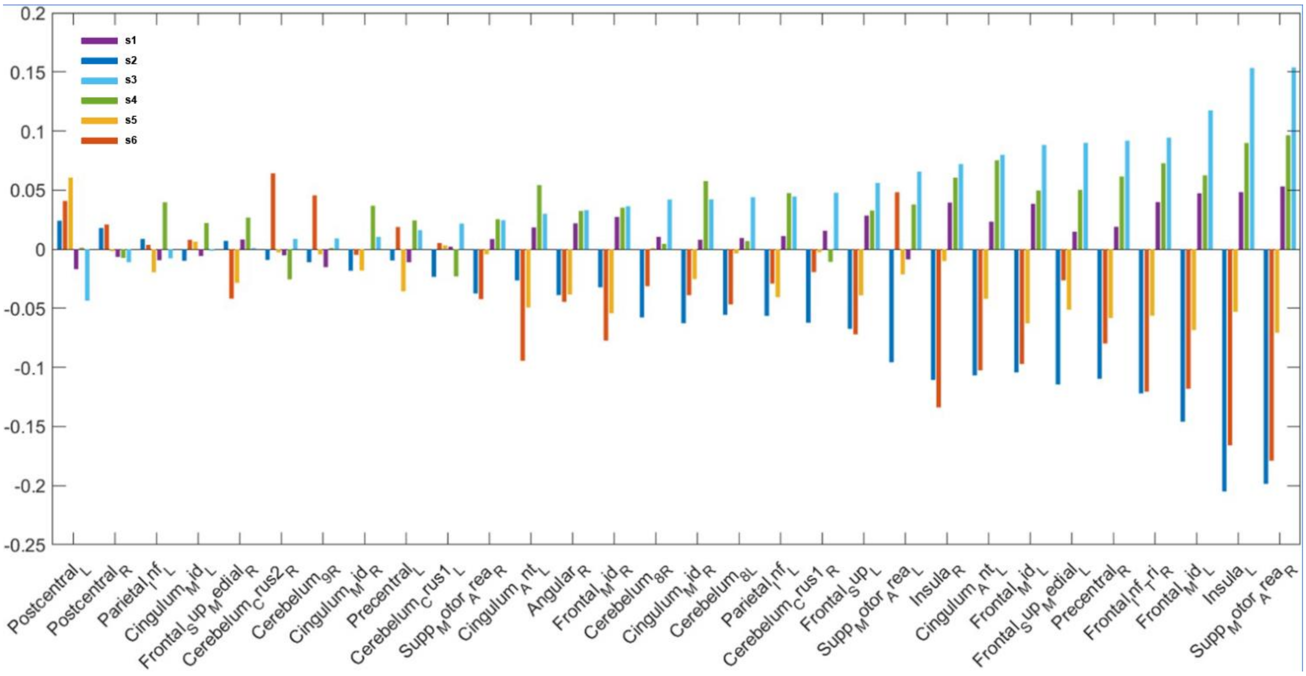


***Figure S4.*** *Brain states' activation patterns in group level.*

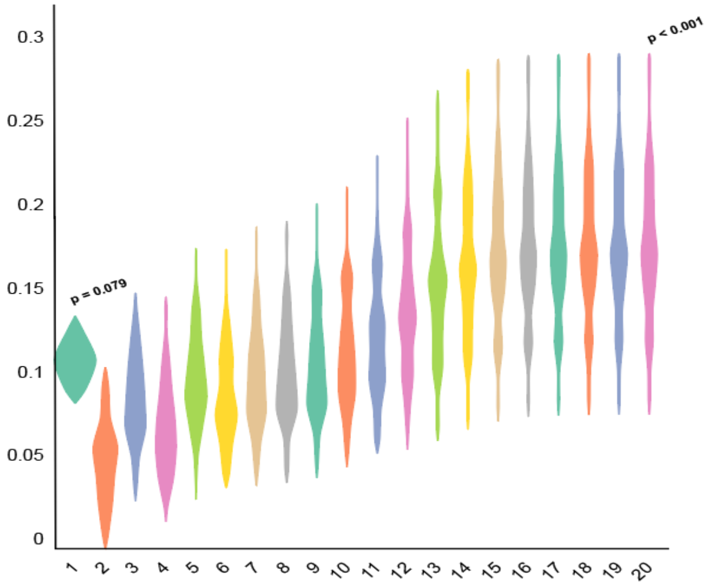


***Figure S5. The adjustments of parameter components in PLSR (Reading).*** *To ensure more reliable results, we applied 10-fold PLSR to randomly divided 305 participants 100 times, adjusting the number of latent components from 1 to 20. The average correlation was used to assess the relationship, with permutation testing (2000 iterations) employed to calculate the p-value. Components number = 20, p-value < 0.001.*

**Supplementary tables**

**Table S1.** Demographics of participants included in the analysis.

| Analysis | n | Males/Females | Age Range | Average Age±SD |
|---|---|---|---|---|
| Math | 310 | 163/147 | 6-12 | 9.151±1.353 |
| Reading | 305 | 159/146 | 6-12 | 9.148±1.342 |

**Table S2.** Brain regions associated with the common neural pattern related to math achievement score across cognitive and affective tasks

| Rank | Node | X | Y | Z | Sum of edge loading | AAL_label |
|---|---|---|---|---|---|---|
| **1** | 30 | 25.2 | 12.4 | 49.4 | 427.1763 | Frontal_Mid_R |
| **2** | 166 | -27.6 | -9.1 | 55.9 | 423.8156 | Precentral_L |
| **3** | 164 | -23.2 | 10.7 | 53.6 | 320.6269 | Frontal_Mid_L |
| **4** | 25 | 7 | -8.1 | 52.9 | 313.189 | Supp_Motor_Area_R |
| **5** | 149 | -39.3 | 17.2 | 46.7 | 240.2537 | Frontal_Mid_L |
| **6** | 161 | -6.5 | -4.3 | 47.6 | 222.5302 | Cingulum_Mid_L |
| **7** | 10 | 8.4 | 53.3 | 23.9 | 211.9306 | Frontal_Sup_Medial_R |
| **8** | 148 | -11.2 | 34.3 | 51.5 | 190.0772 | Frontal_Sup_Medial_L |
| **9** | 158 | -41.6 | -14.7 | 44.8 | 171.9272 | Postcentral_L |
| **10** | 28 | 6.1 | 14 | 48.7 | 171.3094 | Supp_Motor_Area_R |
| **11** | 114 | 23 | -71.8 | -29.1 | 162.9318 | Cerebelum_Crus1_R |
| **12** | 22 | 40 | 17.6 | 29.2 | 159.3485 | Frontal_Inf_Tri_R |
| **13** | 140 | -6 | 48.1 | 11.7 | 146.73 | Cingulum_Ant_L |
| **14** | 84 | 5.3 | -1 | 35.6 | 146.6346 | Cingulum_Mid_R |
| **15** | 253 | -26.3 | -69.5 | -30.6 | 141.5381 | Cerebelum_Crus1_L |
| **16** | 179 | -35.7 | -39.3 | 47.7 | 140.2577 | Parietal_Inf_L |
| **17** | 236 | -6.5 | -66.2 | -37.7 | 139.5478 | Cerebelum_8_L |
| **18** | 162 | -9.1 | 0.4 | 66.5 | 136.5748 | Supp_Motor_Area_L |
| **19** | 100 | 32.2 | -78.5 | -40.4 | 134.8252 | Cerebelum_Crus2_R |
| **20** | 221 | -5.1 | 13.2 | 28.7 | 129.2419 | Cingulum_Ant_L |
| **21** | 146 | -27.3 | 34.1 | 36.4 | 123.6351 | Frontal_Sup_L |
| **22** | 89 | 7.8 | -23.1 | 44.9 | 120.6775 | Cingulum_Mid_R |
| **23** | 48 | 47.8 | -61.6 | 34.7 | 120.1849 | Angular_R |
| **24** | 155 | -32.5 | 22.1 | 5.8 | 120.0185 | Insula_L |
| **25** | 171 | -50.6 | -23.8 | 41.4 | 117.3554 | Parietal_Inf_L |
| **26** | 35 | 41.4 | 3.5 | 7.2 | 117.1255 | Insula_R |
| **27** | 32 | 32 | -5.4 | 52.1 | 112.7194 | Precentral_R |
| **28** | 105 | 6.9 | -67.9 | -37.8 | 111.9052 | Cerebelum_8_R |
| **29** | 33 | 42 | -23.4 | 53.4 | 109.8401 | Postcentral_R |
| **30** | 115 | 7.6 | -56.7 | -50.8 | 108.5254 | Cerebelum_9_R |
| **31** | 91 | 8.3 | -39.9 | 48.1 | 91.96459 | Cingulum_Mid_R |
| **32** | 218 | -7.8 | -22.4 | 46 | 89.88953 | Cingulum_Mid_L |
| **33** | 38 | 32.4 | -39.2 | 49.6 | 88.87557 | Parietal_Inf_R |
| **34** | 85 | 5.1 | -38.9 | 27 | 88.48948 | Cingulum_Post_R |
| **35** | 172 | -23.5 | -31.6 | 63.6 | 86.37543 | Postcentral_L |
| **36** | 254 | -21.2 | -53.4 | -23.6 | 82.93822 | Cerebelum_6_L |
| **37** | 26 | 26.3 | -12.9 | 66.2 | 81.92315 | Precentral_R |
| **38** | 64 | 56.5 | -8.5 | -14.3 | 81.55587 | Temporal_Mid_R |
| **39** | 23 | 57.8 | -8.3 | 27.3 | 78.8887 | Postcentral_R |
| **40** | 225 | -6.5 | -53.9 | 37.4 | 63.02319 | Precuneus_L |
| **41** | 90 | 6.2 | -57.4 | 38.2 | 62.29548 | Precuneus_R |

| | | | | | | |
|---|---|---|---|---|---|---|
| **42** | 145 | -10.1 | 55.7 | 30.2 | 61.34999 | Frontal_Sup_Medial_L |
| **43** | 138 | -6.9 | 48.3 | -5.7 | 60.19294 | Frontal_Med_Orb_L |
| **44** | 226 | -8.8 | -42.6 | 50.1 | 60.01965 | Cingulum_Mid_L |
| **45** | 110 | 21.1 | -54.8 | -23.8 | 58.59388 | Cerebelum_6_R |
| **46** | 242 | -30.2 | -80.2 | -40.3 | 57.31338 | Cerebelum_Crus2_L |
| **47** | 219 | -6 | 34.1 | 26.3 | 56.53693 | Cingulum_Ant_L |
| **48** | 106 | 7.1 | -69 | -20.9 | 53.98631 | Cerebelum_6_R |
| **49** | 141 | -11.7 | 65.1 | 4.2 | 53.09685 | Frontal_Sup_Medial_L |
| **50** | 15 | 6.7 | 21.4 | 31.5 | 53.03531 | Cingulum_Mid_R |
| **51** | 31 | 39.7 | 3.4 | 34 | 52.02413 | Precentral_R |
| **52** | 160 | -16.2 | -19.2 | 69.5 | 51.00494 | Precentral_L |
| **53** | 54 | 50 | -33.8 | -0.7 | 49.21854 | Temporal_Mid_R |
| **54** | 150 | -5 | 17.7 | 46.1 | 46.59127 | Supp_Motor_Area_L |
| **55** | 177 | -28.4 | -62.3 | 40.4 | 32.22945 | Occipital_Mid_L |
| **56** | 5 | 8.2 | 45.9 | -1.7 | 31.54599 | Frontal_Med_Orb_R |
| **57** | 27 | 49.2 | -4.5 | 48.1 | 31.13599 | Precentral_R |
| **58** | 12 | 14.3 | 36.9 | 48.9 | 31.0576 | Frontal_Sup_R |
| **59** | 47 | 54.2 | -45.2 | 36.9 | 30.31876 | SupraMarginal_R |
| **60** | 14 | 40.7 | 14.5 | 48.2 | 30.2455 | Frontal_Mid_R |
| **61** | 40 | 43.3 | -10.8 | 13.9 | 30.2455 | Insula_R |
| **62** | 181 | -59.5 | -25.9 | 21.9 | 29.35149 | SupraMarginal_L |
| **63** | 83 | 7.8 | 34.7 | 17.1 | 29.08937 | Cingulum_Ant_R |
| **64** | 20 | 37 | 20.8 | 5.9 | 28.84863 | Insula_R |
| **65** | 21 | 55.4 | 9.6 | 22.2 | 28.72913 | Frontal_Inf_Oper_R |
| **66** | 231 | -22.7 | -12.8 | -17.4 | 28.18435 | Hippocampus_L |
| **67** | 109 | 23.4 | -59.3 | -52.1 | 28.09828 | Cerebelum_8_R |
| **68** | 157 | -46.2 | 7.9 | 28.6 | 27.00494 | Frontal_Inf_Oper_L |
| **69** | 74 | 45.1 | -74.3 | 2.6 | 26.99012 | Occipital_Mid_R |
| **70** | 53 | 52.8 | 10.9 | -21.8 | 26.67293 | Temporal_Pole_Mid_R |
| **71** | 248 | -8 | -68.4 | -19.9 | 26.15286 | Cerebelum_6_L |
| **72** | 250 | -22.7 | -57.9 | -48.8 | 26.13165 | Cerebelum_8_L |
| **73** | 207 | -25.9 | -63.1 | -12.3 | 26.04618 | Fusiform_L |
| **74** | 147 | -46.1 | 28.2 | 26.8 | 24.82948 | Frontal_Inf_Tri_L |
| **75** | 95 | 28 | -28.4 | -13.7 | 24.7859 | ParaHippocampal_R |
| **76** | 93 | 28.8 | -36.9 | 0 | 24.43264 | Hippocampus_R |
| **77** | 24 | 6 | -22.3 | 65.6 | 23.77718 | Supp_Motor_Area_R |
| **78** | 16 | 53.6 | 24.8 | 0.9 | 23.62194 | Frontal_Inf_Tri_R |
| **79** | 183 | -51.4 | -56.3 | 20.5 | 23.41488 | Temporal_Mid_L |
| **80** | 209 | -48.3 | -67.4 | 1.1 | 23.41488 | Temporal_Mid_L |
| **81** | 144 | -28.8 | 50.1 | 21.7 | 22.8348 | Frontal_Mid_L |
| **82** | 8 | 44.6 | 46.2 | -4.9 | 22.35168 | Frontal_Inf_Orb_R |
| **83** | 167 | -35.9 | -23.3 | 65.6 | 21.7873 | Precentral_L |
| **84** | 56 | 54.5 | -7.7 | -31.5 | 21.69978 | Temporal_Inf_R |
| **85** | 72 | 21 | -63.7 | -9 | 21.69978 | Lingual_R |

Notes: The x, y, and z coordinates were derived from the nodes in the Shen 268 atlas. AAL labels were assigned based on these coordinates using the R package label4MRI (https://github.com/yunshiuan/label4MRI ). Brain regions were ranked by the sum of edge loadings obtained through 2000 bootstraps. The brain regions in the green section represent those appearing before the first turning point in Fig. S3.

**Table S3**. One-way ANOVA results on frequences of brain states in four tasks

| state | F | p-value |
|---|---|---|
| **s1** | 0.92 | 0.519 |
| **s2** | 9.14 | <0.001 |
| **s3** | 4.04 | 0.014 |
| **s4** | 9.42 | <0.001 |
| **s5** | 0.44 | 0.724 |
| **s6** | 2.12 | 0.144 |

Notes: All p-values were corrected using the FDR correction method.

**Table S4**. The correlation between six brain states' frequency and math achievement score

| | States' frequency | | Sample entropy | | Network segregation | | Global efficiency | |
|---|---|---|---|---|---|---|---|---|
| | **r** | **p** | **r** | **p** | **r** | **p** | **r** | **p** |
| **s1** | -0.082 | 0.179 | -0.063 | 0.327 | 0.111 | 0.154 | -0.051 | 0.566 |
| **s2** | 0.227 | <0.001 | 0.214 | <0.001 | 0.151 | 0.048* | -0.031 | 0.710 |
| **s3** | 0.131 | 0.033 | 0.166 | 0.010 | 0.085 | 0.276 | 0.004 | 0.949 |
| **s4** | -0.182 | 0.004 | -0.160 | 0.010 | 0.033 | 0.570 | -0.136 | 0.050 |
| **s5** | -0.007 | 0.902 | -0.004 | 0.945 | 0.040 | 0.570 | -0.084 | 0.287 |
| **s6** | -0.141 | 0.027 | -0.122 | 0.048 | 0.053 | 0.533 | -0.167 | 0.020 |

Notes: All p-values were corrected using the FDR correction method.

*Only 309 participants were included in the calculation of the Spearman correlation between s2 network segregation and math achievement scores for the network segregation algorithm.

**Table S5.** Louvain community detection results on the brain state connectivity pattern

| Region of interest | s1 | s2 | s3 | s4 | s5 | s6` |
|---|---|---|---|---|---|---|
| Frontal_Mid_R | | | | | | |
| Frontal_Mid_L | | | | | | |
| Frontal_Mid_L | | | | | | |
| Frontal_Sup_Medial_R | | | | | | |
| Frontal_Sup_Medial_L | | | | | | |
| Supp_Motor_Area_R | | | | | | |
| Frontal_Inf_Tri_R | | | | | | |
| Cingulum_Ant_L | | | | | | |
| Frontal_Sup_L | | | | | | |
| Angular_R | | | | | | |
| Insula_L | | | | | | |
| Cerebelum_Crus1_R | | | | | | |
| Cerebelum_Crus1_L | | | | | | |
| Cerebelum_8_L | | | | | | |
| Cerebelum_Crus2_R | | | | | | |
| Cerebelum_8_R | | | | | | |
| Cerebelum_9_R | | | | | | |
| Precentral_L | | | | | | |
| Supp_Motor_Area_R | | | | | | |
| Cingulum_Mid_L | | | | | | |
| Postcentral_L | | | | | | |
| Cingulum_Mid_R | | | | | | |
| Parietal_Inf_L | | | | | | |
| Supp_Motor_Area_L | | | | | | |
| Cingulum_Ant_L | | | | | | |
| Cingulum_Mid_R | | | | | | |
| Parietal_Inf_L | | | | | | |
| Insula_R | | | | | | |
| Precentral_R | | | | | | |
| Postcentral_R | | | | | | |

Notes: In a given state, brain regions of the same color were assigned to the same community.
